# Molecular Mechanisms Underlying Human Vγ9Vδ2 T Cell Activation by Butyrophilin-3 (BTN3) Targeted Antibodies

**DOI:** 10.1101/2025.10.21.681109

**Authors:** Bangdong Huang, Weizhi Xin, Wenjia Zhang, Weijie Gao, Yundi Hu, Yuehua Liu, Qiang Su, Qiang Zhou

## Abstract

γδ T cells express T cell receptors (TCRs) composed of paired γ and δ chains. The Vγ9Vδ2 T cells are the main subpopulation of γδ T cells in human peripheral blood, which responds to phosphoantigens (pAgs) through butyrophilins (BTN) molecules, such as BTN3A1/2/3 and BTN2A1. Antibodies targeting BTN3As, such as ICT01, can activate Vγ9Vδ2 T cells independently of pAgs, although the underlying mechanism remains poorly characterized. In this study, we reveal the molecular basis of ICT01-mediated Vγ9Vδ2 T cell activation using structural, biochemical, and cellular analyses. ICT01 binds to a unique region in the extracellular domain of BTN3As, destabilizing the BTN2A1-BTN3As interface and facilitating Vγ9Vδ2 TCR engagement, ultimately resulting in activation of Vγ9Vδ2 T cells. Our findings provide insights into the mechanism by which agonist antibodies induce γδ T cell activation and provide guide strategies for developing BTN-targeted immunotherapies.

## Main

T cells are essential for maintaining immune homeostasis and defending hosts against pathogens^1^. They are divided into αβ and γδ lineages, which express αβ and γδ T cell receptors (TCRs), respectively. γδ T cells are widely distributed across tissues, including skin, mucosal surfaces, liver, and peripheral circulation^2-4^. In humans, γδ T cells only constitute a minor population of circulating T cells, accounting for approximately 2-5%^5^. Unlike αβ T cells, γδ T cells respond to a wide spectrum of non-peptide antigens without MHC restriction^6-8^. The Vγ9Vδ2 T cells are the major γδ T cell subset in the human peripheral blood, which plays a crucial role in antitumor and antimicrobial immunity^9-11^. The Vγ9Vδ2 T cells can be activated by pAgs, such as the endogenous isopentenyl diphosphate (IPP)^12,13^ and the microbial metabolite *E*-4-hydroxy-3-methyl-but-2-enyl diphosphate (HMBPP)^14,15^.

BTN2A1^16-20^ and BTN3A paralogues^20-24^ (including BTN3A1, BTN3A2, and BTN3A3, which share high sequence identity, especially in extracellular region) serve as ligand molecules involving in pAg sensing and Vγ9Vδ2 T cell activation. BTN molecules typically comprise extracellular immunoglobulin variable (IgV) -like and constant (IgC) -like domains, a single transmembrane domain, and a cytoplasmic region containing a juxtamembrane (JM) region and a B30.2 domain (BTN3A2 has no B30.2 domain)^25^. PAg can be bound by the B30.2 domains of BTN3A1 and of BTN2A1 simultaneously, thereby gluing the intracellular regions of BTN2A1 homodimer and BTN3A dimer (the BTN3A1 homodimer, the BTN3A1/BTN3A2 heterodimer, or the BTN3A1/BTN3A3 heterodimer) for TCR engagement^20,24,26-28^. The IgV domain of BTN2A1 interacts with the lateral face of the Vγ9 chain^18,26-28^, whereas the IgV domain of BTN3A (BTN3A1/BTN3A2/BTN3A3) contacts the apical face of both Vδ2 and Vγ9 chains^26-28^. This dual-ligand recognition mechanism stably tethers the Vγ9Vδ2 TCR to the BTN2A1 homodimer and BTN3A homo-/heterodimer, ultimately leading to TCR activation^26-29^.

While pAg-induced Vγ9Vδ2 T cell activation via BTN molecules has been characterized^26-28^, the molecular mechanisms by which anti-BTN3A antibodies modulate Vγ9Vδ2 T cell responses remain unresolved. The agonist monoclonal antibodies (mAbs) 7.2, 20.1, and CTX-2026 can activate γδ T cells independently of pAg^21,30,31^, whereas the 103.2 antibody antagonizes pAg-induced γδ T cell activation in its IgG form^21,30^. The clone of mAb 7.2 elicits degranulation and IFN-γ release at levels comparable to those induced by the well-characterized clone of mAb 20.1^32^. ICT01, a humanized anti-BTN3A antibody derived from mAb 7.2, binds to the IgV domain of BTN3As with sub-nanomolar affinity and triggers specific Vγ9Vδ2 T cell activation, leading to the killing of malignant cells *in vitro*^32^ and showing promising initial safety and tolerability profile in clinical trials^32^.

Prior studies show that all these agonists bind specifically to the IgV domain of BTN3A molecules^30-32^. Although an early model proposed that 20.1 mAb activates Vγ9Vδ2 T cells by inducing multimerization of BTN3A dimers on the cell surface^30^, a recent study demonstrated that such multimerization is dispensable for signaling^33^. Notably, multiple studies have indicated that BTN3A1 alone is insufficient to elicit Vγ9Vδ2 T cell activation upon stimulation^16,17,19,34^, and the agonistic activity of mAb 20.1 requires the presence of BTN2A1^16,17^. Furthermore, emerging studies have revealed that BTN2A1 and BTN3A molecules can form complex^26-28^. These findings highlight the functional interdependence of BTN2A1 and BTN3A in γδ T cell activation and underscore the necessity of re-examining the molecular mechanisms wherein anti-BTN3A antibodies such as ICT01 modulate the interaction between the BTN2A1–BTN3A complex and the Vγ9Vδ2 TCR.

In this study, we solved cryo-EM structures of ICT01 Fab bound to BTN molecules, including the BTN2A1/BTN3A1/BTN3A2–ICT01 complex and the BTN3A1/BTN3A2–ICT01 complex. We propose that ICT01 binding to BTN3A extracellular regions triggers the dissociation of the BTN2A1/BTN3A oligomeric complex, enabling the Vγ9Vδ2 TCR to engage the IgV domains of both BTN2A1 and BTN3A in the dual-ligand mode, ultimately leading to the activation of Vγ9Vδ2 T cells. Our study reveals the structural mechanism underlying agonistic antibody-mediated activation of Vγ9Vδ2 T cells and provides a blueprint for developing BTN-targeting immunotherapies.

## Results

### ICT01 Fab interacts with the BTN2A1/BTN3A1/BTN3A2 complex

To investigate the interaction of ICT01 with BTN molecules, we purified full-length BTN2A1/BTN3A1/BTN3A2 complex, as well as the Fab fragment of ICT01 (Fig. 1a; Supplementary Fig. S1a,b). SDS-PAGE analysis showed that the BTN2A1, BTN3A1, and BTN3A2 proteins co-migrated, indicating the formation of a stable complex (abbreviated as ‘BTN complex’), consistent with recent studies^26-28^. ICT01 Fab was then incubated with the BTN complex either in the presence or absence of HMBPP, followed by SEC analysis. Under both conditions, ICT01 co-migrated with the BTN complex, demonstrating that ICT01 binds to the BTN complex independently of HMBPP (Supplementary Fig. S1c,d).

**Fig. 1.**
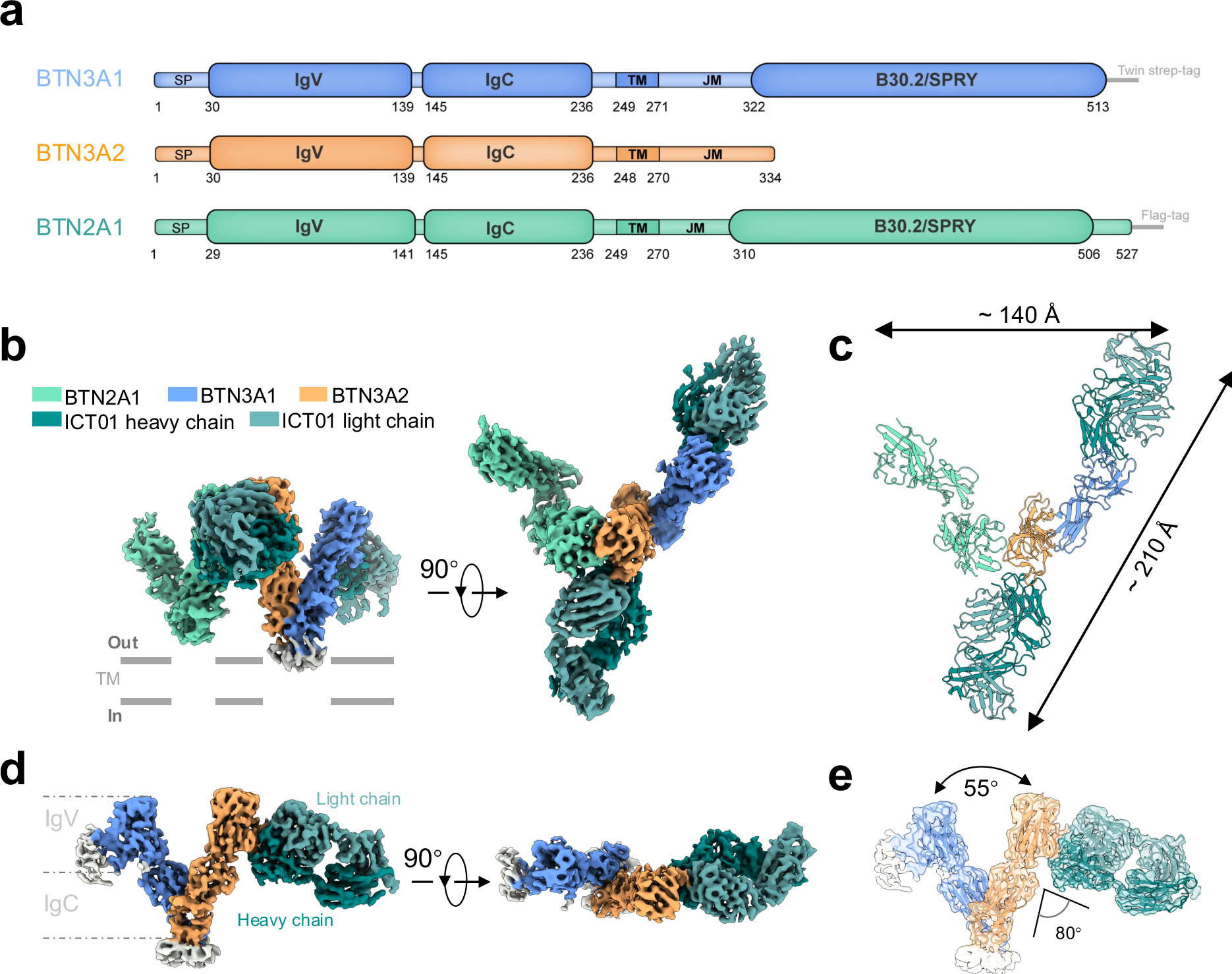
Structures of ICT01 Fab-bound BTN complexes. **a** Domain architectures and construct designs of BTN molecules. **b-c** Cryo-EM map (**b**) and atomic model (**c**) of the BTN2A1/BTN3A1/BTN3A2 complex bound by two ICT01 Fabs. The map, shown in two orthogonal views, is contoured at 7.8 σ. **d-e** Cryo-EM map (**d**) and superposition of model and map (**e**) of the BTN3A1/BTN3A2 heterodimer bound by a ICT01 Fab. The map is contoured at 7.1σ. Subunits are colored as follows: BTN2A1 (medium aquamarine), BTN3A1 (cornflower blue), BTN3A2 (sandy brown), BTN3A3 (khaki), ICT01-H (teal) and ICT01-L (cadet blue). This color scheme is consistently applied throughout the study.

In the absence of HMBPP, the SEC profile of the ICT01–BTN complex shows two distinct peaks, designated ‘Peak 1’ and ‘Peak 2’ (Supplementary Fig. S1d). SDS-PAGE analysis indicated that Peak 1 fractions contained all the components including BTN2A1, BTN3A1, BTN3A2, and ICT01, whereas Peak 2 fractions showed reduced levels of BTN2A1 (Supplementary Fig. S1d).

### Overall structures of the ICT01-BTN complex

To explore the detailed interaction between ICT01 and BTN molecules, we separately pooled and concentrated the fractions from Peak 1 and Peak 2 for cryo-electron microscopy (cryo-EM) analysis, separately. From the Peak 1 sample, we successfully determined the cryo-EM structures of the BTN2A1/BTN3A1/BTN3A2–ICT01 complex at 3.6 Å resolution and the BTN3A1/BTN3A2–ICT01 complex at 3.7 Å resolution, respectively (Supplementary Fig. S2 and S3). Further several rounds of 3D classification of the BTN2A1/BTN3A1/BTN3A2–ICT01 complex yielded a 4.0 Å resolution structure containing two ICT01 Fabs bound to the BTN complex (Fig. 1b,c; Supplementary Fig. S2 and S3). The cytoplasmic regions, comprising the juxtamembrane (JM) domains and B30.2 domains, remained unsolved, suggesting a high degree of conformational flexibility (Supplementary Fig. S2b). In contrast, recent studies revealed that pAg binding rigidifies the JM and B30.2 segments, enabling their structural determination^26-28^. The 2D averages of Peak 2 confirmed that two ICT01 Fabs bind to ectodomain of BTN3A1 and BTN3A2, respectively (Supplementary Fig. S4a,b). However, preferred particle orientation impeded the reconstruction of the BTN3A1/BTN3A2–ICT01 complex. Therefore, all subsequent structural analyses of the BTN3A1/BTN3A2–ICT01 complex were based on data derived from Peak 1 sample.

The ectodomains of the BTN2A1 homodimer or the BTN3A1/BTN3A2 heterodimer form “V-shaped” architectures mediated by intra-dimer IgC-IgC interactions, respectively (Fig. 1b,c; Supplementary Fig. S5a-c). These dimers associate in a 1:1 stoichiometric ratio to form a “W-shaped” tetramer (“BTN tetramer”), buttressed by head-to-head packing of the CFG faces (β-strands C″, C′, C, F and G) from the BTN2A1 and BTN3A2 IgV domains (Fig. 1c; Supplementary Fig. S5b). One ICT01 Fab binds to the IgV domain of BTN3A2 near the BTN2A1 IgV domain, while the other interacts with the IgV domain of BTN3A1 within the BTN tetramer, spanning a distance of approximately 210 Å (Fig. 1c). The overall structure of BTN3A1/BTN3A2–ICT01 complex closely resembles those observed within the BTN2A1/BTN3A1/BTN3A2-ICT01 complex (Fig. 1d,e; Supplementary Fig. S5e), with a root mean square deviation (RMSD) of 1.1 Å.

### Interaction of ICT01 with the ectodomain of the BTN complex

All structurally characterized agonist antibodies targeting the CFG face of BTN3A IgV domain are spatially proximal to Vγ9Vδ2 TCR (Supplementary Fig. S6a). Structural analysis reveals that ICT01 adopts a binding mode analogous to the 20.1^30^ and Th001^27^ antibodies, with their epitopes near the epitope of the CTX-2026^31^ antibody (Supplementary Fig. S6b,c). The epitopes of those four agonist antibodies are located adjacent to or only minimally overlapping with the Vγ9Vδ2 TCR footprint on BTN3A (Supplementary Fig. S6c). ICT01 antibody can bind to BTN3A1 or BTN3A2 molecules (Fig. 2b,d). These two binding interfaces are very similar due to the nearly identical amino acid sequences of the IgV domains (Supplementary Fig. S7). The following structural analysis takes the binding interface between ICT01 antibody and BTN3A2 as an example. The ICT01 Fab binds the BTN3A2 IgV domain at about 80° docking angle, mediated primarily by the heavy chain complementarity-determining region (CDR) 1–3 and light chain CDR1 and CDR3 (Fig. 2a). The heavy chain CDR3, which is the longest CDR of 15 residue length, extends its loop directly into a relatively hydrophilic pocket near the IgV^BTN2A1^-IgV^BTN3A1^ interface (Supplementary Fig. S5f).

**Fig. 2.**
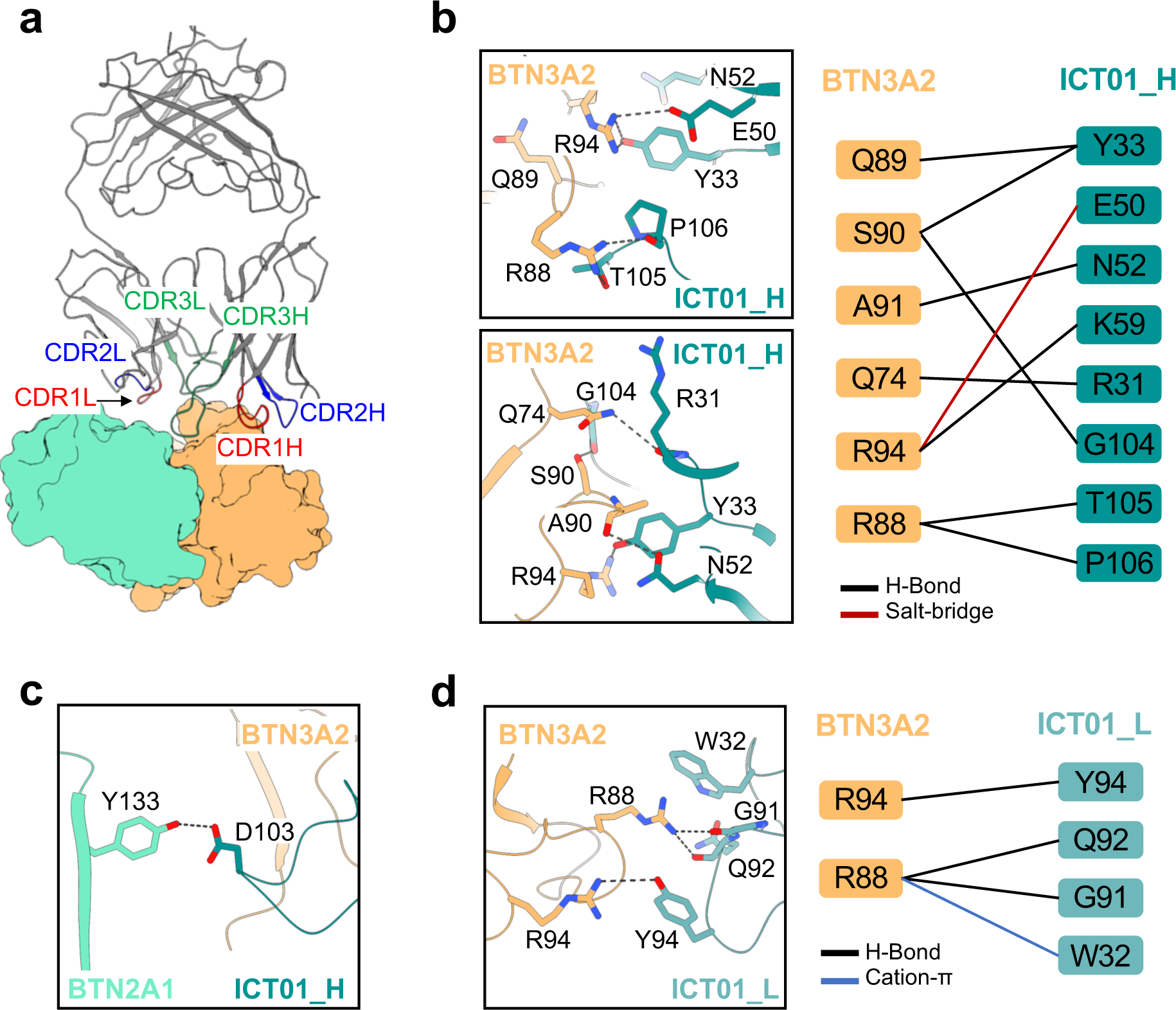
Interactions between ICT01 and ectodomains of BTN complex. **a** The CDR1-3H and CDR1-2L regions of ICT01 interact with the BTN complex. ICT01 is displayed in a cartoon representation, while the IgV^BTN2A1^ and IgV^BTN3A2^ are shown as surface style. Other regions are omitted for clarity. **b** Close-up views of the interface between IgV^BTN3A2^ and ICT01 heavy chain (left panel). Key interactions are illustrated by a residue contact map (right panel). **c** Close-up view of the hydrogen bond at the interface between IgV^BTN2A1^ and ICT01-H. **d** Close-up views of the interface between IgV^BTN3A2^ and ICT01 light chain (left panel). Key interactions are depicted by a residue contact map (right panel).

ICT01 buries approximately 642.3 Å² area on the BTN3A2 surface in the BTN2A1/BTN3A1/BTN3A2–ICT01complex. Detailed inspection of the IgV^BTN3A2^–ICT01 interfaces reveals an intricate interaction network that comprises at least 11 hydrogen bonds, a cation-π interaction, a salt bridge interaction, and extensive van der Waals interactions. The side chain groups of Y33^ICT01-H^ and T105^ICT01^ form hydrogen bonds with R94^BTN3A2^ and R88^BTN3A2^, respectively, while the side chain of E50^ICT01-H^ forms a salt-bridge with R94^BTN3A2^ (Fig. 2b; Supplementary Fig. S8a,b). Additionally, we also observed a hydrogen bond between D103^ICT01H^ and Y133^BTN2A1^ (Fig. 2c; Supplementary Fig. S8c), which was first remarked for the anti-BTN3A antibody. The light chain features a cation-π interaction between W32^ICT01-L^ and R88^BTN3A2^, accompanied by hydrogen bonds formed between G91^ICT01-L^ and R88^BTN3A2^, Q92^ICT01-L^ and R88^BTN3A2^, and Y94^ICT01-L^ and R94^BTN3A2^ (Fig. 2d; Supplementary Fig. S8d). R94^BTN3A^ serves as a key interaction hub, contacted by both the heavy and light chains of ICT01 (Fig. 2b,d). Meanwhile, the ICT01 epitope on BTN3A does not encompass residues V68 and R73, which are critical for pAg signaling^18^, the latter of which is also essential for direct TCR Vδ2 chain binding^26-28^.

### ICT01 binding destabilizes the BTN2A1–BTN3A interaction

To explore the working mechanism of ICT01, we compared the structures of the BTN2A1/BTN3A1/BTN3A2–ICT01 complex and the BTN3A1/BTN3A2–ICT01 complex. Conformational changes in BTN3A2 IgV domain (loop 70-73aa) is revealed, despite the fact that residues such as R73^BTN3A2^ does not specifically interacts with ICT01 (Fig. 2b). In the BTN2A1/BTN3A1/BTN3A2–ICT01 complex, R73^BTN3A2^ forms a cation-π interaction with F71^BTN2A1^ and establishes a salt bridge with E135^BTN2A1^, while Y127^BTN3A2^ also interacts with F71^BTN2A1^ (Fig. 3a; Supplementary Fig. S8e). In contrast, within the BTN3A1/BTN3A2–ICT01 complex, R73^BTN3A2^ forms an intrachain hydrogen bond with its own Y127 ^BTN3A2^ (Fig. 3b; Supplementary Fig. S8f). Similar interaction patterns have been observed in agonist-bound BTN3A1 structures, such as those bound to 20.1^30^ (PDB ID: 4F9L) and CTX-2026^31^ (PDB ID: 6XLQ) (Fig. 3c; Supplementary Fig. S6a,b).

**Fig. 3.**
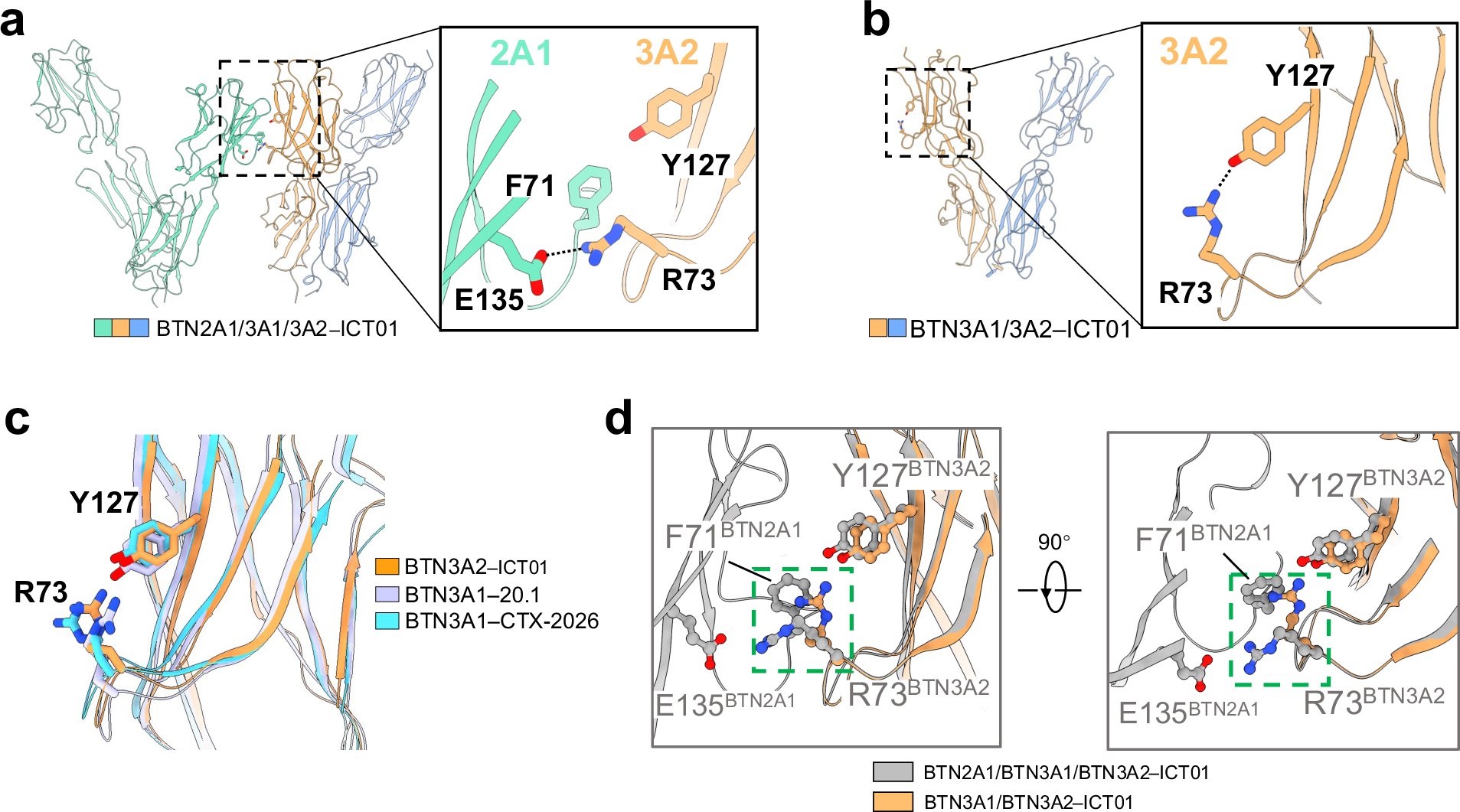
ICT01 binding destabilizes BTN2A1–BTN3A interactions. **a** R73^BTN3A2^ forms a salt bridge with E135^BTN2A1^ and a cation–π interaction with F71^BTN2A1^ within the IgV^BTN2A1^– IgV^BTN3A2^ interface. **b** R73^BTN3A2^ forms an intrachain hydrogen bond with Y127^BTN3A2^ in the ICT01–BTN3A1/BTN3A2 complex. **c** A similar intrachain hydrogen bond is observed in 20.1– bound BTN3A1 and CTX-2026–bound BTN3A1. **d** Superposition of the BTN3A1/BTN3A2– ICT01 structure onto the BTN2A1/BTN3A1/BTN3A2–ICT01 structure reveals steric clash between the BTN3A1 within the ICT01–bound BTN3A1/3A2 and BTN2A1 within the ICT01– bound BTN2A1/BTN3A1/BTN3A2 structure. Structures of antibody Fabs are omitted for clarity in panels **a–d**.

Structural alignment based on the BTN3A2 IgV domain reveals that if R73^BTN3A2^ adopts a conformation similar to that seen in the BTN3A1/BTN3A2–ICT01 complex, it would clash sterically with F71^BTN2A1^ at the IgV^BTN2A1^–IgV^BTN3A2^ interface within the BTN2A1/BTN3A1/BTN3A2–ICT01 complex (Fig. 3d). These findings suggests that ICT01 binding can disrupt the IgV^BTN2A1^–IgV^BTN3A2^ interaction. Furthermore, mutation of R73^BTN3A1^ or Y127^BTN3A1^ weakens the association between BTN2A1 and BTN3A1 on the cell surface^18^, highlighting the critical role of these residues in maintaining the structure of the BTN2A1/BTN3A1/BTN3A2 complex.

### ICT01 promotes TCR engagement by disassembling the BTN oligomer

Previous work showed that BTN2A1 and BTN3A1 homodimers can assemble into higher-order oligomers via constitutive IgV domain interactions^18^ (Supplementary Fig. S9a). Given the near-identical amino acid sequences of BTN3A1 and BTN3A2 IgV domains (Supplementary Fig. S8), the BTN2A1/BTN3A1/BTN3A2 complex are capable of forming oligomeric structures, which is supported by findings from our other study^28^.

Structural comparisons show nearly identical V-shaped angles among BTN3A dimers across both BTN2A1/BTN3A1/BTN3A2–ICT01 and BTN3A1/BTN3A2–ICT01 complexes. However, the BTN2A1 homodimer and the BTN3A1/BTN3A2 heterodimer in the BTN2A1/BTN3A1/BTN3A2–ICT01 complex exhibit 3° and 8° wider angles, respectively, compared to those BTN2A1 homodimer and BTN3A1 homodimer in the BTN2A1/BTN3A1 complex crystal structure (Fig. 4a). These expanded angles resemble those seen in the HMBPP-bound BTN2A1/BTN3A1/BTN3A2 tetramer^28^. Superposition reveals that these increased angles would cause steric clashes with adjacent BTN molecules in the context of the higher-order oligomer (Supplementary Fig. S9b). Therefore, we hypothesize that the ICT01 binding to BTN3As within the oligomer disrupts the BTN2A1-BTN3A interaction, leading to the dissociation of oligomer into tetramers or tetramer into dimers.

**Fig. 4.**
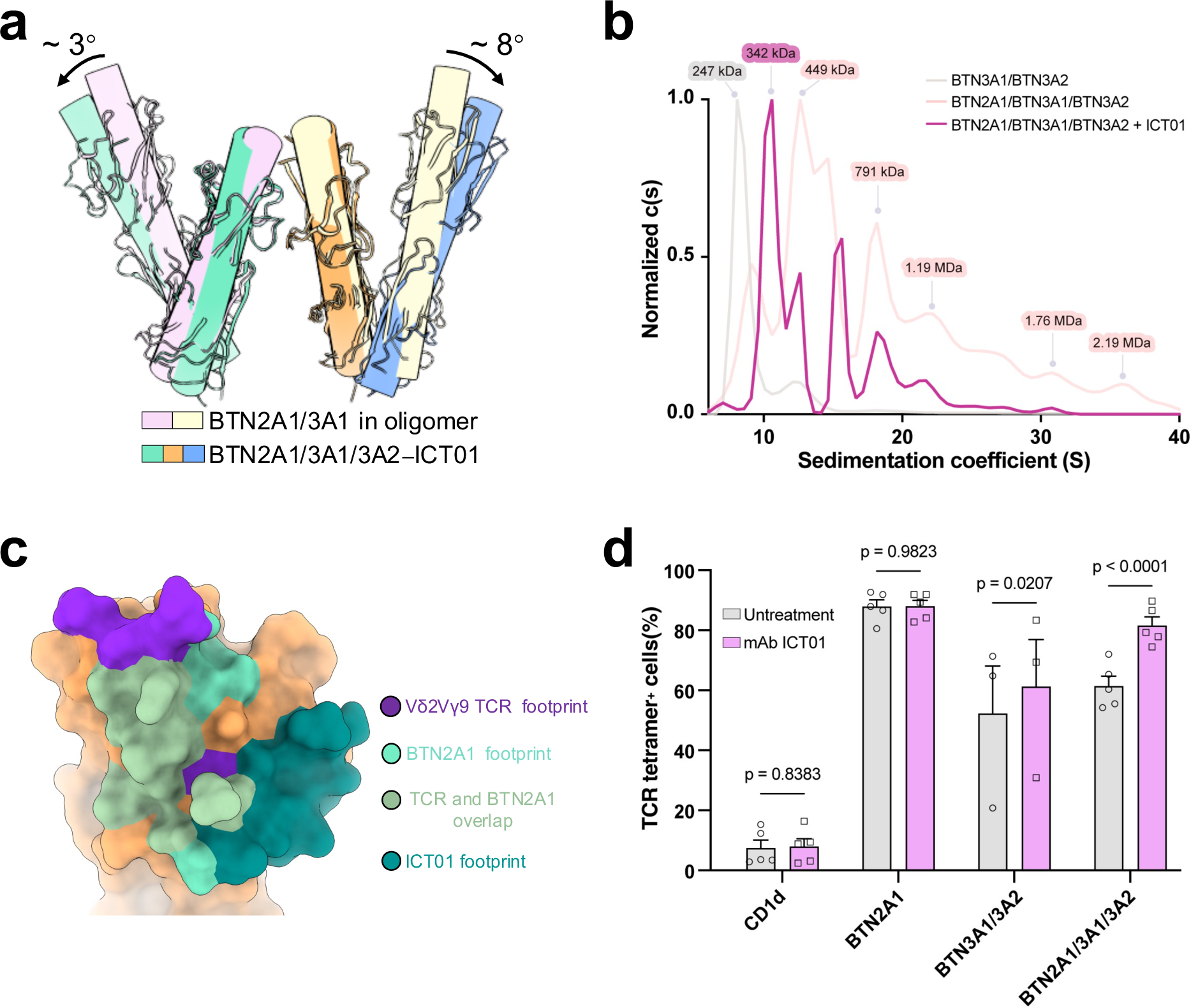
Conformation changes upon ICT01 binding enhance TCR engagement with the BTN complex. **a** Superposition of the BTN2A1/BTN3A1/BTN3A2 tetramer from ICT01-BTN2A1/BTN3A1/BTN3A2 complex with the crystal structure of the BTN2A1/BTN3A1 ectodomain complex. The V-shaped angles of the BTN2A1 dimer and BTN3A1/BTN3A2 heterodimer are larger than those of the BTN2A1 dimer and BTN3A1 dimer in the crystal structure of the BTN2A1/BTN3A1 ectodomain complex. **b** Analytical ultracentrifugation (AUC) sedimentation profiles of the BTN3A1/BTN3A2 complex and BTN2A1/BTN3A1/BTN3A2 complex (± ICT01). ICT01 treatment induces dissociation of high-mass oligomers into lower-mass species. Raw data were normalized to 0–1. **c** Mapping of BTN2A1 binding sites, Vγ9Vδ2 TCR binding sites, and ICT01 epitopes on BTN3A2. The BTN3A2 is shown as a surface representation. **d** G115 TCR tetramer staining of NIH-3T3 fibroblasts expressing CD1d, BTN2A1, BTN3A1/BTN3A2, and BTN2A1/BTN3A1/BTN3A2 in the presence or absence of ICT01. *P* value was determined by two-way ANOVA with Dunnett’s multiple comparison test. Bar graphs depict the mean ± s.e.m. Data are pooled from four independent experiments, except for BTN2A1/BTN3A1, which was analyzed in three independent experiments.

We performed analytical ultracentrifugation (AUC) analysis to monitor changes of the molecular weight of the BTN2A1/BTN3A1/BTN3A2 complex upon addition of ICT01. The profile of BTN2A1/BTN3A1/BTN3A2 alone shows high levels of higher molecular weight species (HMWS), corresponding to the oligomeric assemblies of the complex (Fig. 4b). Upon addition of ICT01, these HMWS are markedly diminished, with a concomitant increase in lower-molecular-weight species (Fig. 4b). This shift in sedimentation profile suggesting that ICT01 binding triggers the disassembly of the BTN oligomer into smaller tetrameric and/or dimeric units.

BTN2A1 and BTN3A molecules occlude each other’s Vγ9Vδ2 TCR binding sites (Fig. 4c; Supplementary Fig. S9c), and their compact organization within oligomers may further restrict TCR access (Supplementary Fig. S9d). Vγ9Vδ2 TCR tetramer staining revealed that ICT01 pretreatment did not impair TCR binding to cells expressing BTN2A1. Despite the spatial proximity between the ICT01 epitope and the Vγ9Vδ2 TCR binding site on BTN3A (Fig. 4c), pretreatment with ICT01 does not interfere with TCR staining cells expressing BTN3A1/BTN3A2. ICT01 treatment significantly enhanced TCR tetramer binding to cells co-expressing the BTN2A1/BTN3A1/BTN3A2 complex (Fig. 4d), which forms higher-order oligomers on the plasma membrane^28^. Together, these findings demonstrate that ICT01-induced dissociation of the oligomer into tetramers enhances the efficacy of Vγ9Vδ2 TCR engagement (Fig. 4d).

## Discussion

BTN2A1 and BTN3As are ubiquitously expressed in immune cells^35-37^. To allow for rapid and precise sensing of pAg levels during pathogen invasion or cellular stress, while preventing aberrant activation of γδ T cells under normal conditions, cells must employ a finely tuned regulatory mechanism to maintain immune homeostasis. In our other work, we found BTN2A1 and BTN3As form higher-order oligomers and proposed a pAg-driven BTN oligomer disassembling model for the pAg-induced Vγ9Vδ2 T cell activation^28^. BTN molecules assemble into oligomers that sterically occlude Vγ9Vδ2 TCR engagement, thereby preventing inappropriate activation. PAg binding induces bending of extracellular regions and stabilizes the intracellular regions of BTN molecules, inducing steric clash in the intracellular regions and promoting dissociation of the oligomers into tetrameric units that are primed for TCR engagement^28^.

Agonistic antibodies such as ICT01 disassemble the oligomer in a pAg-independent mode by directly targeting the IgV domain of BTN3A. Binding of the agonistic antibody ICT01 induces conformational changes that disrupt the inter-chain interactions between BTN2A1 and BTN3As, leading to the disassembly of the BTN2A1/BTN3As oligomer into tetrameric and/or dimeric units that subsequently engage the Vγ9Vδ2 TCR in the dual-ligand mode to activate T cells (Fig. 5). Our findings indicate that both pAg-induced and agonist antibody-induced Vγ9Vδ2 T cell activation require structural disassembly of the BTN2A1/BTN3As oligomer, wherein destabilization of interchain interactions permits TCR engagement, thus working in a unified model.

**Fig. 5.**
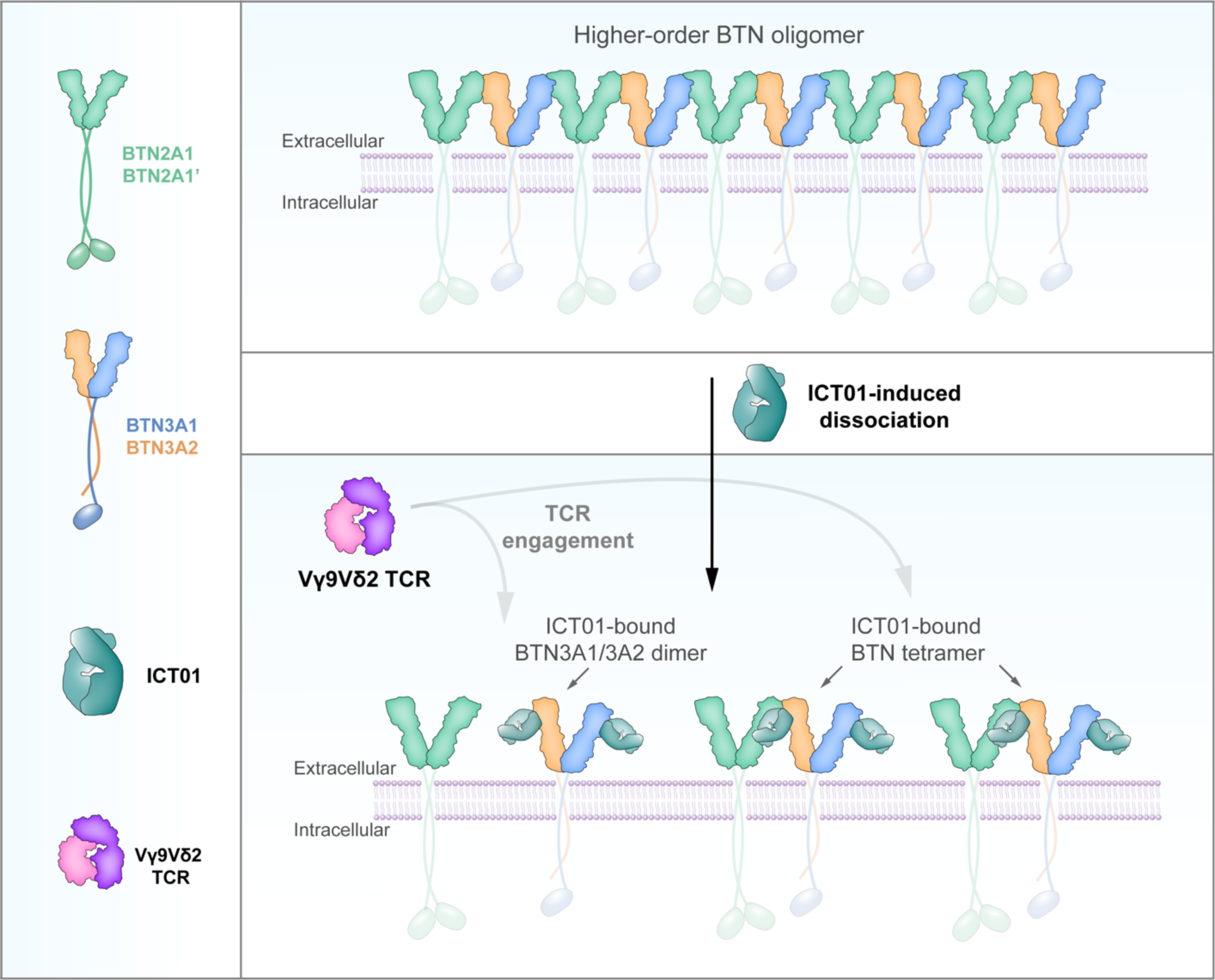
A schematic diagram of the activation mechanism of ICT01. In the absence of pAgs or agonist, the BTN2A1 homodimer and BTN3A heterodimer assemble into higher-order oligomers through ectodomain interactions (upper panel). The intracellular regions of these oligomer exhibit conformational flexibility, depicted by transparent cartoons (upper panel). This oligomeric architecture sterically precludes Vγ9Vδ2 TCR engagement by maintaining tight intermolecular interactions between BTN molecules, thereby occluding access of Vγ9Vδ2 TCR to the IgV^BTN2A1^-IgV^BTN3A2^ interface. Binding of ICT01 to the IgV domains of BTN3A induces conformational changes in the extracellular domains, triggering dissociation of the oligomer into tetrameric and/or dimeric units that can engage the Vγ9Vδ2 TCR (bottom panel).

Within the BTN2A1/BTN3A1/BTN3A2-ICT01 complex, ICT01 interacts with BTN2A1 through a hydrogen bond between D103^ICT01-H^ and Y133^BTN2A1^ (Fig. 2c; Supplementary Fig. S8c). The interaction of the BTN3A-targeted antibody with BTN2A1 also exists in the recently reported BTN2A1/BTN3A1/BTN3A2-Th001 complex^26^. This finding suggests that antibody engineering should consider the potential impact of interactions with BTN2A1.

Notably, ICT01 and other agonist antibodies bind BTN3A IgV domain without obstructing or with slightly overlapping with the TCR interaction sites, thereby enabling TCR engagement following oligomer dissociation. On the other hand, agonist antibodies need to disrupt BTN2A1/BTN3A oligomers, so their epitopes need to be close to the interface of BTN2A1/BTN3A. This can explain why the epitopes of these agonist antibodies are very close to each other, which represents a privileged antigenic patch for development of agonist antibodies. With γδ T cell-based therapeutics advancing rapidly^38^, our findings not only provide novel insights into the mechanism of agonistic antibodies but also offer a conceptual framework for the development of immunotherapies.

## Materials and Methods

### Construct design

The cDNAs encoding human BTN2A1, BTN3A1, and BTN3A2 were codon-optimized and synthesized by Tsingke Biotechnology Co., Ltd. These cDNAs were subsequently subcloned into an optimized pCAG vector. DNA encoding anti-BTN3A antibody heavy and light chains (ICT01), codon-optimized for expression in ExpiHEK293F cells, was cloned into separate pCAG plasmids. A synthetic signal peptide (MDMRVPAQLLGLLLLWLSGARC) was fused to the N-terminus of BTN2A1, BTN3A1, BTN3A2, and the light chain of ICT01, while another signal peptide (MEFGLSWLFLVAILKGVQC) was fused to the N-terminus of the heavy chain of ICT01. To facilitate protein purification, a Flag tag (MDYKDDDDK) was inserted at the C-terminus of BTN2A1 and BTN3A2, while a twin-strep II tag (WSHPQFEKGGGSGGGSGGSAWSHPQFEK) was added to the C-terminus of BTN3A1. Additionally, an 8×His tag was inserted at the C-terminus of the constant region of the ICT01 heavy chain.

### Expression and purification of BTN2A1/BTN3A1/BTN3A2 complex

For protein expression and purification of BTN2A1/BTN3A1/BTN3A2 complex, 2.4 mg of plasmids encoding BTN2A1, BTN3A1, and BTN3A2 (0.8 mg each) were pre-incubated with 5 mg of 40K polyethyleneimine (PEI, Yeasen) in 40 mL of fresh medium (Sino Biological Inc.) for 30 minutes. The mixture was transfected into 0.8 liters of ExpiHEK293F cells (Thermo Fisher Scientific Inc.), which were then cultured for 3 days at 37°C with 5% CO2 in a Multitron-Pro shaker (Infors, 120 rpm) before collection. ExpiHEK293F cells expressing BTN2A1/BTN3A1/BTN3A2 complex was centrifuged at 4000g for 10 min. The cells were then resuspended into 25 mM HEPES (pH 7.4), 150 mM NaCl, 2 μg/mL pepstatin, 1 μg/mL aprotinin, and 2 μg/mL leupeptin. The cell membranes were solubilized overnight at 4°C using 1% (w/v) LMNG and 0.1% CHS (Anatrace). Soluble fraction was isolated by centrifugation at 12,500 rpm for 1 hour, and pass over 2mL of anti-Flag M2 affinity resin (MilliporeSigma), washed with wash buffer (25 mM HEPES, pH 7.4, 150 mM NaCl, 0.02% (w/v) GDN), and eluted using the wash buffer supplemented with 400 μg/mL Flag peptide. The eluate from the anti-Flag resin was loaded onto Strep-Tactin XT resin (IBA), washed with the wash buffer, and eluted using a buffer containing 25 mM HEPES (pH 7.4), 150 mM NaCl, 0.02% (w/v) GDN, and 50 mM D-biotin. Finally, the eluted complex was concentrated to 0.8 mL and further purified using size-exclusion chromatography (SEC) on a Superose 6 Increase 10/300 column (GE Healthcare).

### Expression and purification of ICT01 Fab

ExpiHEK293F cells expressing the Fab format of ICT01 antibody were cultured at 37°C under 5% CO_2_ for 5 days before harvesting. After centrifuging at 4000g for 20 minutes, the supernatant was collected and filtered through a 0.22 μm membrane to remove cell debris. The filtrate was then concentrated to approximately 150 mL and the fab format of ICT01 antibody was affinity purified by nickel-chelation chromatography (Qiagen). The resin was washed with a buffer containing 25 mM HEPES (pH 7.4), 150 mM NaCl, and 30 mM imidazole. The ICT01 Fab was eluted using the wash buffer supplemented with 300 mM imidazole. The resulting eluate was concentrated and further purified via SEC employing a Superdex 200 Increase 10/300 GL column in a buffer containing 25 mM HEPES (pH 7.4) and 150 mM NaCl.

### Preparation of the BTN2A1/BTN3A1/BTN3A2–ICT01 Fab complex

The BTN2A1/BTN3A1/BTN3A2 complex was incubated with an excess molar ratio of the ICT01 Fab at 4 °C for 1 hour, either without or with 40 µM HMBPP. The mixture was then concentrated to a final volume of 0.8 mL. Excessive ICT01 Fab was subsequently removed using SEC (Superose 6 increase 10/300, GE Healthcare) in a buffer containing 25 mM HEPES (pH 7.4), 150 mM NaCl, 0.02% (w/v) GDN, and 4 µM HMBPP. Finally, the purified TCR-BTN complex was concentrated and prepared for cryo-EM sample.

### Analytical Ultracentrifugation (AUC)

Sedimentation velocity analytical ultracentrifugation (AUC) was conducted using an Optima AUC A/I analytical ultracentrifuge (Beckman Coulter) at 20 °C. The protein complex was diluted to a final volume of 380 μL in AUC buffer (25 mM HEPES, pH 7.4, 150 mM NaCl, 0.02% GDN), and samples were loaded into double-sector sapphire cells alongside 400 μL of reference solution (25 mM HEPES, pH 7.4, 150 mM NaCl). Data were acquired using absorbance detection at 280 nm with a 50 Ti rotor spinning at 40,000 rpm. Extinction coefficients for the protein samples were calculated using the ExPASy ProtParam tool (https://www.expasy.org). Molecular weights and sedimentation coefficients (S) were determined using SEDFIT.

### Construction of NIH3T3 stable cell lines

To generate NIH3T3 stable cell lines expressing distinct BTN variants, the BTN2A1/BTN3A1/BTN3A2 complex linked by P2A peptides or BTN3A1/BTN3A2, BTN2A1, along with an mCherry tag, were cloned into the lentiCRISPR v2 vector under the control of the spleen focus-forming virus (SFFV) promoter. The lentiCRISPR plasmids were co-transfected with pMD2.G and psPAX2 into Lenti-X 293T cells to produce lentiviral particles. Viral supernatants were collected at 48- and 72-hours post-transfection. Lentiviruses were concentrated using 80 μg/mL polybrene (Macklin) and 80 μg/mL chondroitin sulfate C (Macklin), then used to transduce NIH3T3 cells. Transduced cells exhibiting comparable mCherry fluorescence intensity were isolated by fluorescence-activated cell sorting.

### Tetramerization of Soluble Vγ9Vδ2 TCR and Flow Cytometry Staining

Soluble Vγ9Vδ2 TCR containing an Avi tag was enzymatically biotinylated using recombinant BirA biotin ligase, prepared in-house. Biotinylated TCR was incubated with Streptavidin–FITC (Solarbio) at a 4:1 molar ratio overnight at 4 °C to form the Vγ9Vδ2 TCR tetramer complex. NIH3T3 cells expressing BTN2A1, BTN3A1/BTN3A2, or BTN2A1/BTN3A1/BTN3A2 were seeded in 24-well plates and were then harvested using 0.25% trypsin-EDTA, washed three times with ice-cold FACS buffer (PBS containing 2% FBS). The cells were treated with or without ICT01 for 1 h at 4 °C and subsequently incubated with 5 μg of the TCR tetramer complex at 4 °C for 1 h in the dark. After washing, cells were resuspended in FACS buffer and analyzed on a CytoFLEX LX-5L2 flow cytometer (Beckman Coulter). Data were processed using CytExpert software.

### Cryo-EM sample preparation and data collection

To increase the hydrophilicity of the surface of the grid, holey carbon grids (Quantifoil, Au, 300- mesh, R1.2/1.3) were glow-discharged at 15 mA for 30 s. A 3-3.5 μL aliquot of the protein sample was applied to the grid, which was then blotted for 3 s under 100% humidity at 8°C. The grid was then rapidly plunged into liquid ethane using a Vitrobot Mark IV (Thermo Fisher Scientific).

Grids were then transferred to a Krios electron microscopy (Thermo Fisher Scientific) operating at 300 kV, equipped with a Gatan K3 Summit detector and a GIF Quantum energy filter. EPU (Thermo Fisher Scientific) was utilized for automated movie collection. Movie stacks were acquired with a defocus range from -1.0 µm to -2.0 µm in super-resolution mode. The magnification was 81,000X, corresponding to a physical pixel size of 1.087 Å per pixel. Each movie was recorded for 2.56 s, with a total dose of 50 e^-^/Å² distributed across 32 frames.

### Cryo-EM data processing

Movies stacks were initially motion-corrected using MotionCor2 v1.2.6^39^, and dose-weighted micrographs were subsequently imported into cryoSPARC v4^40^. Particles were picked through template-based picking. The particles were extracted and binned by a factor of 4, then subjected to multiple rounds of 2D classification and 2D selection. This process was repeated with the binning factor progressively reduced from 4 to 2, and finally to 1. The raw particles were subject to triple Ab-Initio Reconstruction. There were two classes that exhibited significant protein features. Class I demonstrated dimer binding to ICT01 Fab, whereas Class V exhibited tetramer binding to ICT01 Fab. The particles were initially subjected to NU-refinement and then transferred to RELION5^41^ for 3D classification (skip alignment) and 3D refinement (with mask), ultimately yielding high-quality reconstructions of Class I and Class V at 3.7 Å resolution each. To further improve the local resolution, we conducted consensus refinement using a soft mask that included only the IgV domain and ICT01 Fab signals. This approach yielded local reconstructions of Class I and Class V at 3.7 Å and 3.5 Å resolution, respectively. To reconstruct the complete density of the tetramer bound to two Fabs, the center of the particles was shifted to the corresponding BTN3A1 IgV-Fab interface. The recentered particles were then re-extracted and subjected to NU-refinement, resulting in a reconstruction at 3.8 Å resolution, with the density of the two Fabs clearly visible. Then, this portion of particles was transferred to RELION-5.0^41^ (https://github.com/3dem/relion/tree/ver5.0) using the csparc2star.py script from the UCSF pyem^42^ program packages. A spherical mask was applied to aid in several rounds of RELION 3D classification (excluding alignment), following which, particles from the two distinct classes, exhibiting clear Fab characteristics, were combined after duplicates were removed. These particles were then subjected to RELION 3D refinement, resulting in a reconstruction at 4.0 Å resolution. Subsequently, the particles were subjected to consensus refinement, with the densities of the constant region of Fabs masked out, ultimately yielding a reconstruction at 4.0 Å resolution.

The resolution of the reconstruction was determined by the gold-standard Fourier shell correlation (FSC) using the FSC = 0.143 criterion, either in cryoSPARC v4^40^ or RELION-5^41^ (https://github.com/3dem/relion/tree/ver5.0). All figures depicting the maps and models were prepared using UCSF ChimeraX^43^.

### Model Building

The density maps were sharpened using B-factors. The atomic model of ICT01 Fab, predicted by AlphaFold3^44^, and the atomic model of BTN2A1/BTN3A1/BTN3A2 (PDB ID: <u>8ZHR</u>) were utilized as initial models. The model was rigidly docked into the density map using UCSF Chimera^45^, and then the resulting model was flexibly docked into the same density map using NAMDINATOR^46^ (https://namdinator.au.dk/namdinator/). The fitting model then underwent several iterative rounds of manual adjustment in COOT^47^ and Phenix^48^ real-space refinement, with the incorporation of secondary structure and geometric restraints. To monitor overfitting, we conduct cross-validation between the model and the corresponding maps. In short, the resultant model was refined against one of the two half maps and then computed the FSC curves with both of the two half maps. Also, the resultant model was refined against the summed map of the two half maps and computed the FSC curve. The minor differences among the three curves indicate that the model has not overfitted to the noise. This suggests the reliability of the resultant model and density maps. All three FSC curves were plotted in GraphPad Prism. All models in this work were validated using the Phenix.validation_cryoem^48^ tool or the online server at http://molprobity.biochem.duke.edu. Detailed statistics for the cryo-EM data collection, 3D reconstruction, model refinement and validation are presented in Supplementary Table S1.

## Data availability

The atomic coordinates for all the protein complexes mentioned above have been deposited in the Protein Data Bank under the accession numbers 9KWZ (BTN2A1/BTN3A1/BTN3A2**–**ICT01, one Fab), 9L1P (BTN2A1/BTN3A1/BTN3A2**–**ICT01, one Fab, local refinement), 9KWE (BTN3A1/BTN3A2**–**ICT01), 9L1O (BTN3A1/BTN3A2**–**ICT01, local refinement), 9L1Q (BTN2A1/BTN3A1/BTN3A2-ICT01, two Fabs).

For the BTN2A1/BTN3A1/BTN3A2**–**ICT01 (one Fab) complex, the 3D cryo-EM maps have been deposited in the Electron Microscopy Data Bank under the accession numbers <u>EMD-62622</u> and <u>EMD-62751</u> (one Fab, local refinement). For the BTN2A1/BTN3A1/BTN3A2**–**ICT01 (two Fabs) complex, the 3D cryo-EM maps have been deposited in the Electron Microscopy Data Bank under the accession numbers <u>EMD-62752</u>. For the BTN3A1/BTN3A2-ICT01 complex, the 3D cryo-EM maps have been deposited in the Electron Microscopy Data Bank under the accession numbers <u>EMD-62608</u> and <u>EMD-62750</u> (local refinement). Previously published structures referred to in this paper include: <u>8ZHR</u>. All data are available in the manuscript and the Supplementary Information. Source data are provided with this paper. All materials are available from the corresponding authors upon reasonable request.

## Acknowledgements

This work is supported by National Key R&D Program of China (2020YFA0509300) and the Key Regional Research and Development Program (2023CSJZN0600) from Ministry of Science and Technology of China, Natural Science Foundation of Zhejiang Province (QKWL25C0501), National Natural Science Foundation of China (324B2025, 82241081), State Key Laboratory of Gene Expression, and Shenzhen Medical Academy of Research and Translation. We thank the cryo-EM facility, the high-performance computing center, the protein characterization and crystallography facility of Westlake University for technical assistance, advice, and support. We thank Dr. Q. Huang for the generous gift of Lenti-X 293T.

## Author contributions

Q.Z., B.H., and W.X. conceived the project and designed the experiments. W.X. conducted cloning and protein purification. B.H. performed cryo-EM analysis. W.X. perform biochemical assay and other cell-based assay. All authors contributed to data analysis. Q.Z., B.H., and W.X. wrote the manuscript. Q.Z. supervised the project.

## Supplementary figures and tables

**Supplementary Fig. S1.**
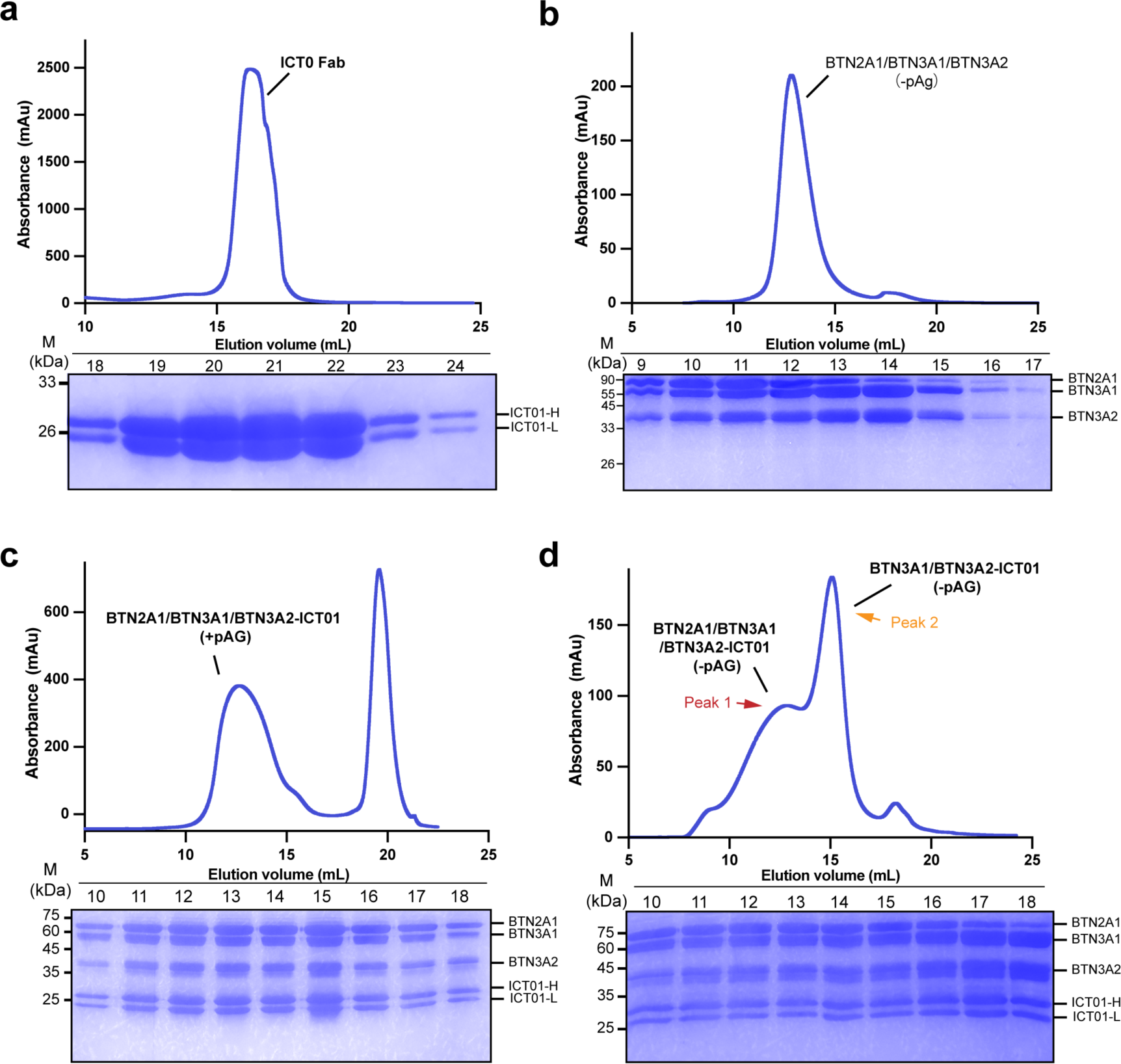
Biochemical characterization of ICT01 Fab and BTN complex. **a** Representative size exclusion chromatography (SEC) profile of the ICT01 Fab (top panel) and analysis of the peak fractions by Coomassie blue staining after SDS-PAGE (bottom panel). **b** Representative SEC profile of the BTN complex (top panel) and SDS-PAGE analysis of the peak fractions (bottom panel). **c** Representative SEC profile of the ICT01-Fab and BTN complex incubated with HMBPP (top panel) and corresponding SDS-PAGE analysis (bottom panel). **d** Representative SEC profile of the ICT01-Fab and BTN complex incubated without HMBPP (top panel) and SDS-PAGE analysis (bottom panel). The components with different molecular weights are labeled on the right side of the gel in the bottom panels in **c** and **e**. ICT01-H: the heavy chain of the ICT01 Fab fragment; ICT01-L: the light chain of the ICT01 Fab fragment;

**Supplementary Fig. S2.**
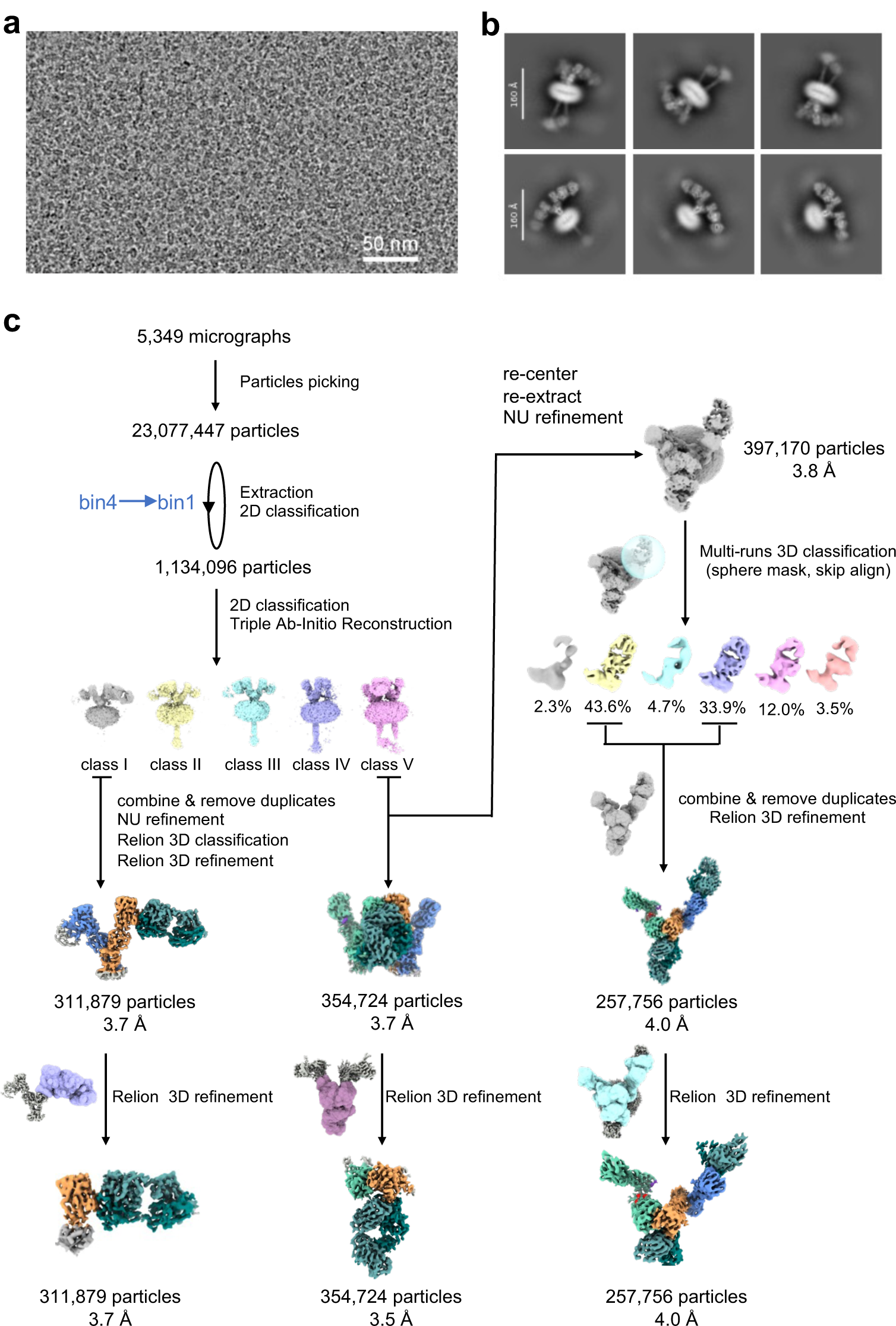
Cryo-EM analysis of ‘Peak 1’ sample. a-b. Representative cryo-EM micrograph (**a**) and 2D class averages (**b**) of the ‘Peak 1’ dataset. The 2D class averages reveal that two ICT01 Fabs bind to the ectodomain of the BTN2A1/BTN3A1/BTN3A2 tetramer complex and the BTN3A1/BTN3A2 heterodimer complex. **c** Cryo-EM data processing workflow for the ‘Peak 1’ dataset. Details are described in Methods section.

**Supplementary Fig. S3.**
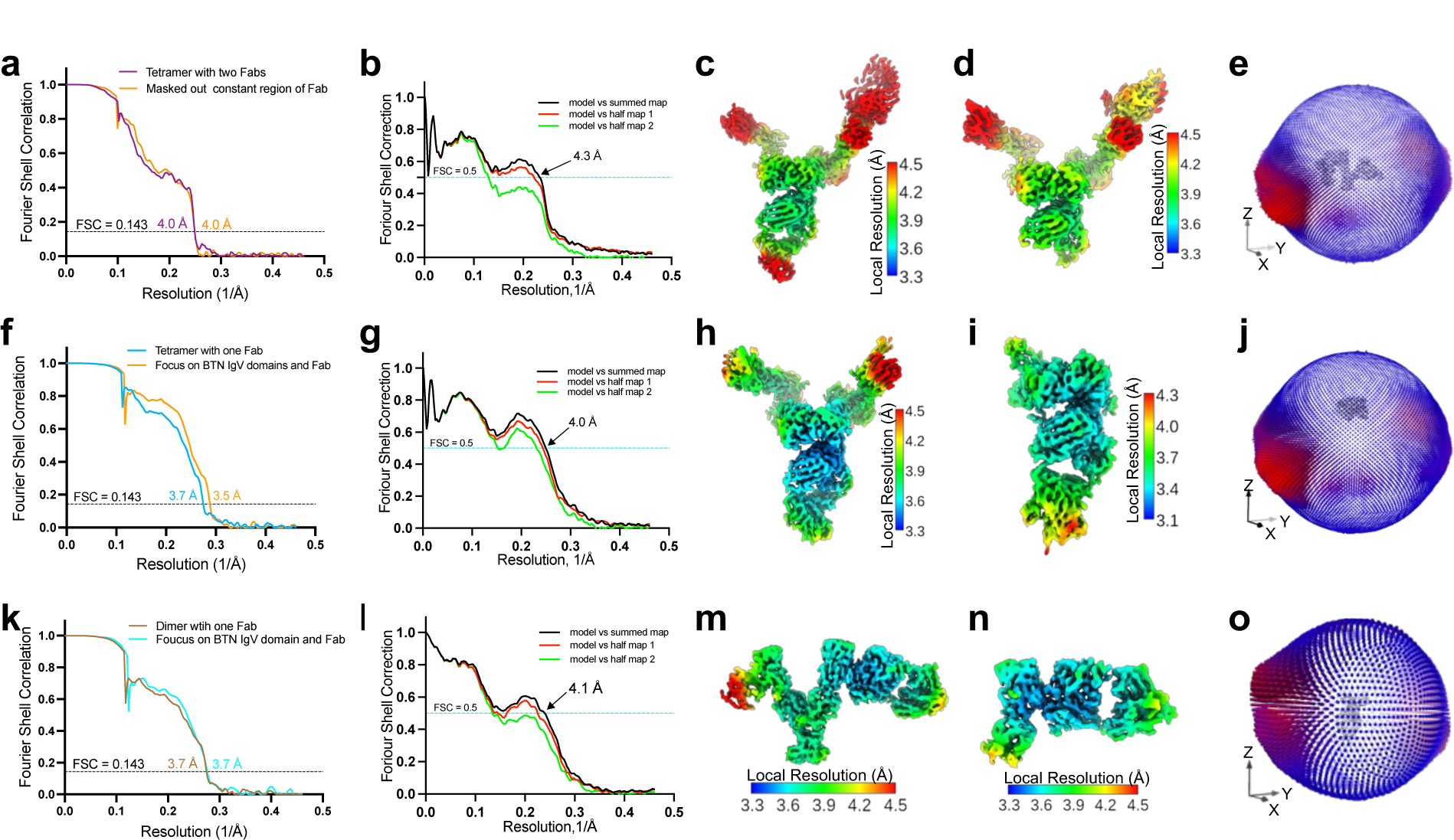
Validation of cryo-EM maps and models of ICT01-BTN2A1/BTN3A1/BTN3A2 complex and ICT01-BTN3A1/BTN3A2 complex. **a** Fourier shell correlation (FSC) curves for the global ectodomain refinement and local refinement of ICT01-BTN2A1/BTN3A1/BTN3A2 complex (two ICT01 Fabs). **b** Model versus map FSC curves for the overall ICT01-BTN2A1/BTN3A1/BTN3A2 complex (two ICT01 Fabs). **c-d** Local resolution estimation of the global ectodomain refinement map and local refinement map. **e** Angular distribution of particles used in the final refinement. **f** Fourier shell correlation (FSC) curves for the global ectodomain refinement and local refinement of ICT01-BTN2A1/BTN3A1/BTN3A2 complex (one ICT01 Fab). **g** Model versus map FSC curves for the overall ICT01-BTN2A1/BTN3A1/BTN3A2 complex (one ICT01 Fab). **h-i** Local resolution estimation of the global ectodomain refinement and local refinement of ICT01-BTN2A1/BTN3A1/BTN3A2 complex (one ICT01 Fab). **j** Angular distribution of particles used in the final refinement. **k** Fourier shell correlation (FSC) curves for the global ectodomain refinement and local refinement of the ICT01-BTN3A1/BTN3A2 complex. **l** Model versus map FSC curves. **m-n** Local resolution estimation of the global ectodomain refinement and local refinement of ICT01-BTN3A1/BTN3A2 complex. **o** Angular distribution of particles used in the final refinement. Details and explanations for the three curves in map versus model plot are provided in the Methods section.

**Supplementary Fig. S4.**
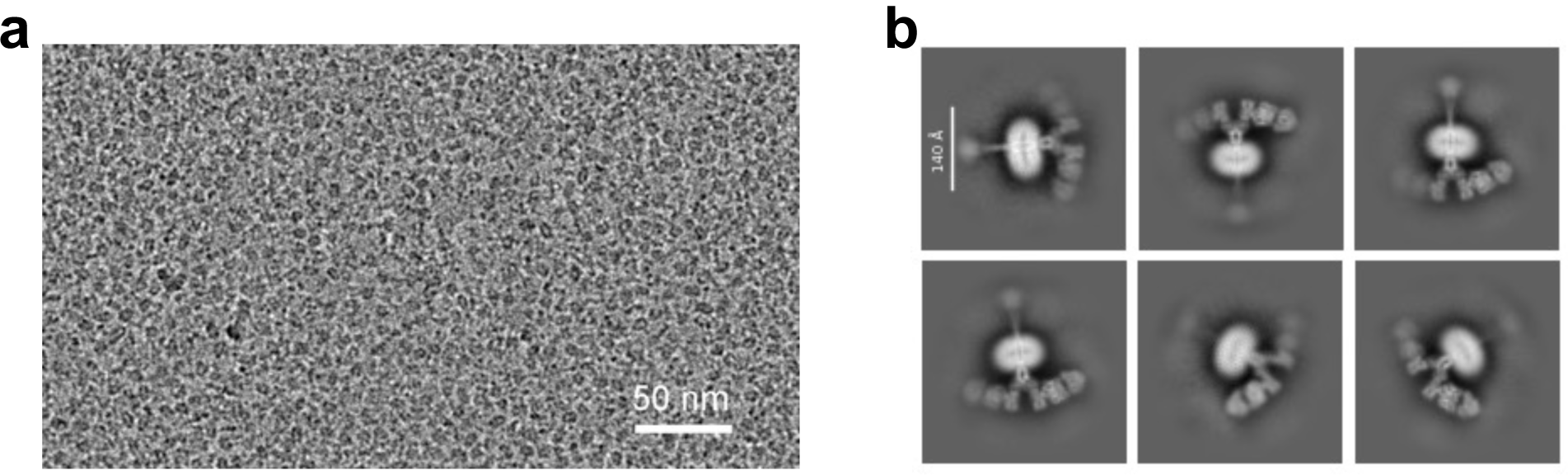
Cryo-EM analysis of ‘Peak 2’ samples. **a-b**, Representative cryo-EM micrograph **(a)** and 2D class averages **(b)** of the ‘Peak 2’ dataset. The 2D class averages reveal that two ICT01 Fabs interact with the ectodomain of the BTN3A1/BTN3A2 heterodimer complex.

**Supplementary Fig. S5.**
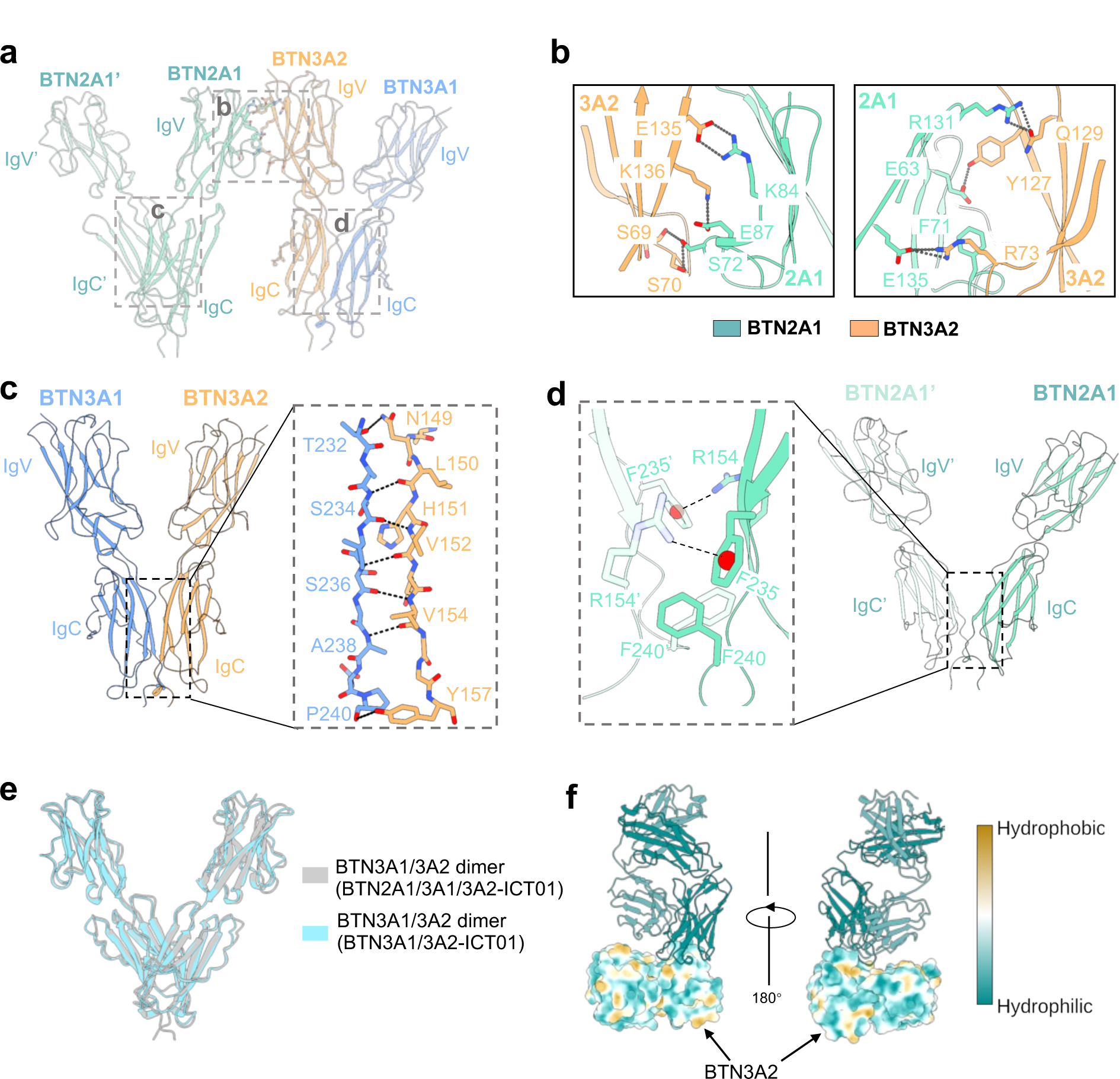
Interchain interactions between BTN molecules. **a** The IgV–IgV and IgC–IgC interchain interactions stabilize the tetrameric assembly. **b** Close-up views of the interface between the IgV domains of BTN2A1 dimer and BTN3A1/BTN3A2 heterodimer. **c-d** IgC domain interactions of the BTN3A1/BTN3A2 heterodimer (**c**) and the BTN2A1 homodimer (**d**) within the tetramer. **e** The V-shaped BTN3A1/BTN3A2 dimer is nearly identical in both the ICT01-bound BTN2A1/BTN3A1/BTN3A2 complex and the ICT01-bound BTN3A1/BTN3A2 complex. **f** Hydrophobic and hydrophilic analysis of the IgV domains of BTN2A1 and BTN3A2. The ICT01 Fab is shown in cartoon representation.

**Supplementary Fig. S6.**
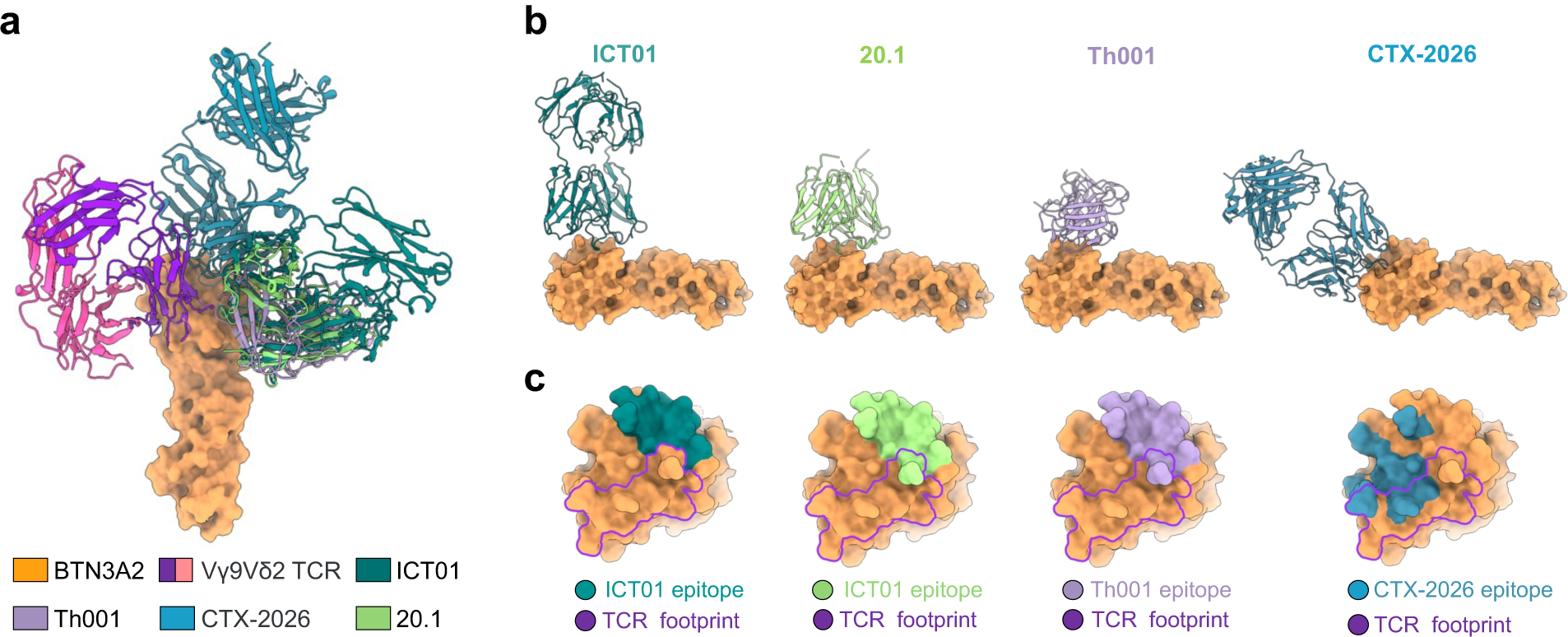
Comparison of binding mode and epitopes between ICT01 and other reported agonists of BTN3As. **a** Structural alignment of BTN3A2 in complex with agonists or the Vγ9Vδ2 TCR. BTN3A2 is depicted as surface, while agonists and the TCR are shown as ribbon cartoons. **b** Comparation of the binding mode of each agonist. **c** Comparison of agonist epitopes with the TCR binding footprint^28^ on BTN3A2. Agonist epitopes are represented as filled shapes, whereas the TCR binding footprint is delineated by a purple line. The epitopes are defined as the residues on BTN3A located within a distance of less than 4 Å from the bound agonist antibody using ChimeraX^43^.

**Supplementary Fig. S7.**
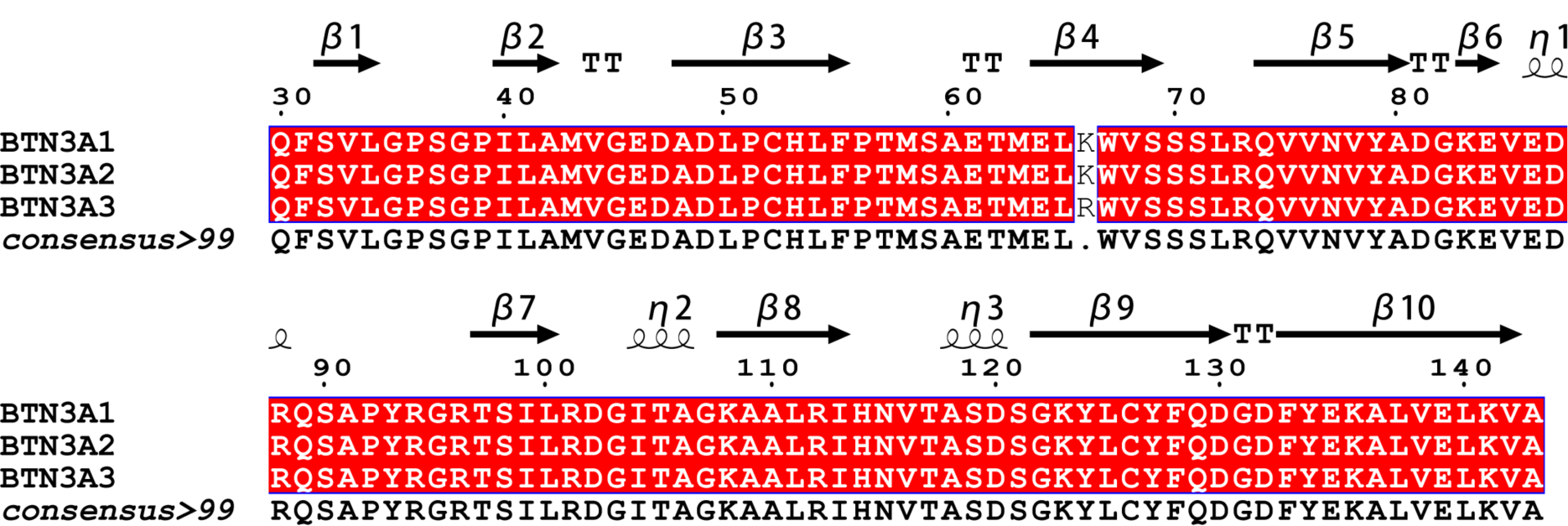
Amino acid sequence alignment of the IgV domains of BTN3A isoforms. Amino acid sequences of the IgV domains (residues 30–143) of BTN3A1 (Uniprot accession: O00481), BTN3A2 (Uniprot accession: P78410), and BTN3A3 (Uniprot accession: O00478) were downloaded from UniProt, with the boundaries of the IgV domain defined by InterPro.

**Supplementary Fig. S8.**
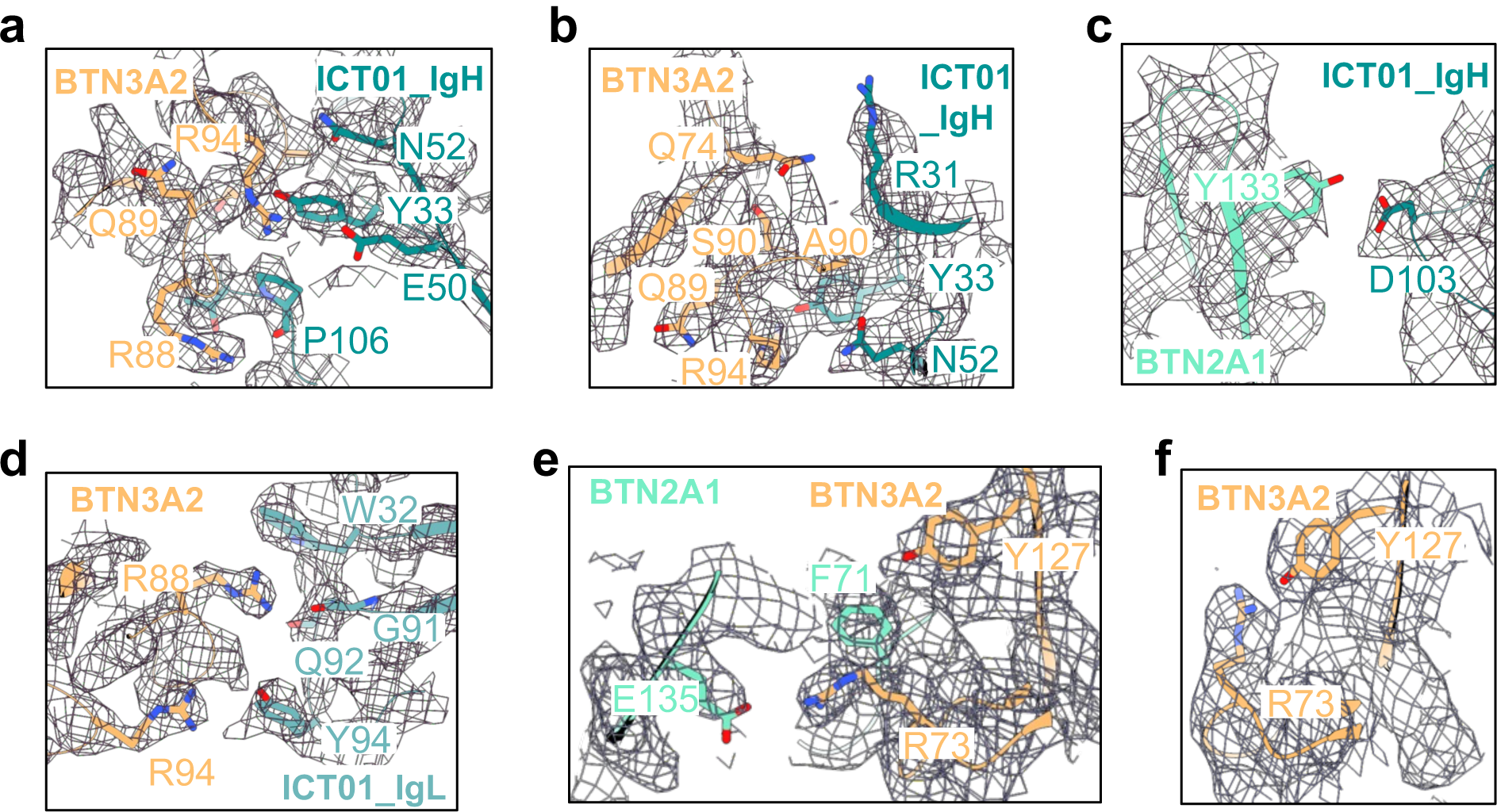
Representative cryo-EM maps of interchain and intrachain interfaces. a-b. Close-up views of the interface between ICT01 Fab and IgV^BTN3A2^, with the cryo-EM map contoured at 10.3 σ. **c** Close-up view of interface between ICT01 heavy chain and IgV^BTN2A1^, with the map contoured at 9.3 σ. **d** Close-up view of the interface between IgV^BTN2A1^ and IgV^BTN3A2^, with the map contoured at 10.6 σ. **e** Close-up view of the intrachain interaction between residues R73 and Y127 within the BTN3A2 IgV domain. The map is contoured at 8.4 σ.

**Supplementary Fig. S9.**
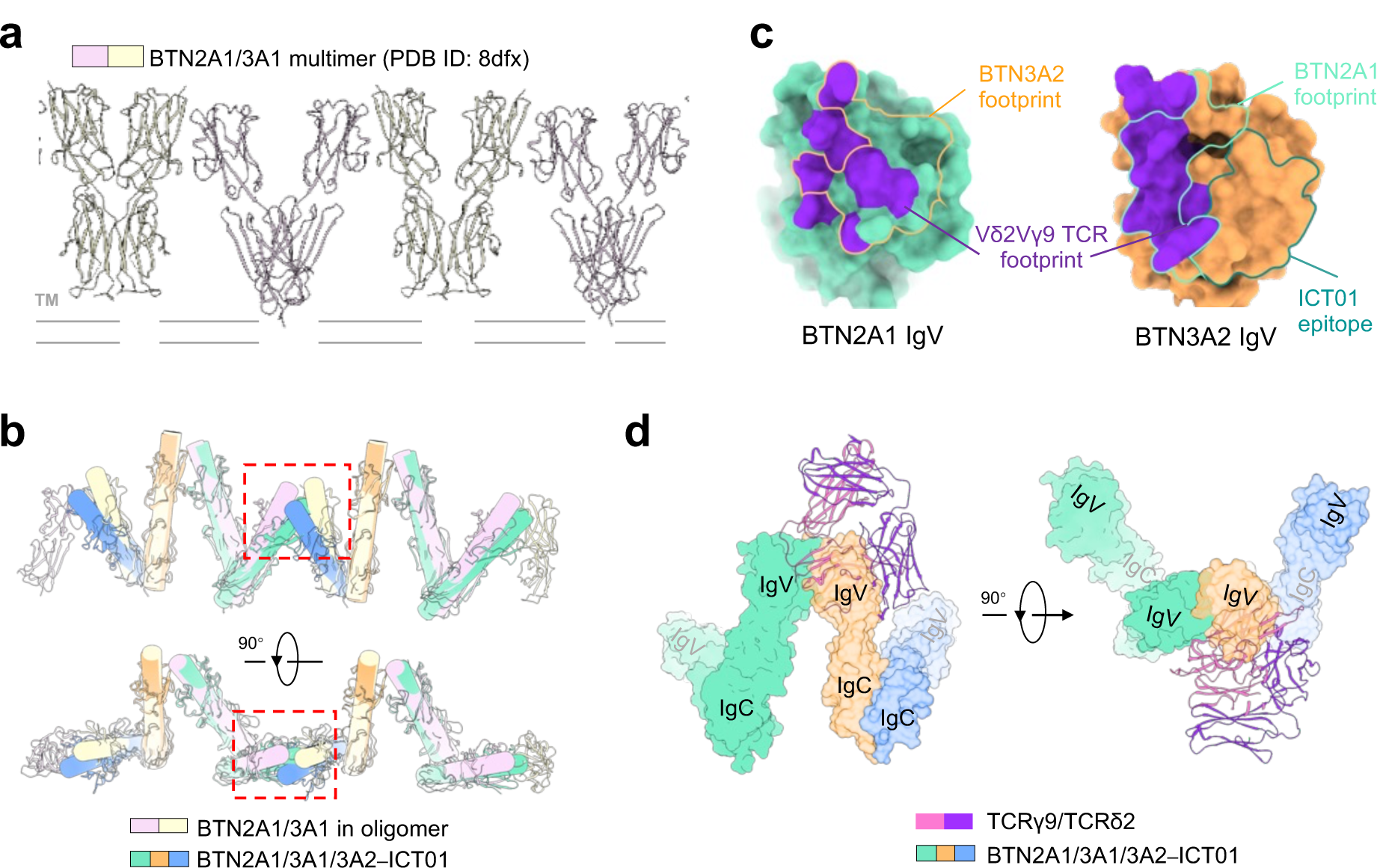
Agonistic antibody binding to BTN3A induces conformation change of BTN oligomer. **a** The linear oligomer of the BTN2A1 and BTN3A1 homodimer (PDB: 8DFX). **b** Superposition of two ICT01 bound–BTN2A1/BTN3A1/BTN3A2 complexes onto the BTN2A1/BTN3A1 oligomer based on the IgV^BTN2A1^–IgV^BTN3A1^ interface. The larger angle of V-shaped dimers poses a substantial challenge in terms of steric hindrance within the adjacent BTN molecules, as indicated by the red dashed box. **c** Footprint mapping of the Vγ9Vδ2 TCR on the IgV domain of BTN2A1 and BN3A2 and of epitope on the BTN3A2 IgV domain. The unfilled shapes highlight the BTN3A2 footprint on the IgV domain of BTN2A1, as well as the BTN2A1 footprint and ICT01 footprints on the IgV domain of BTN3A2, as indicated. **d** Superposition of the Vγ9Vδ2 TCR–BTN2A1 complex onto the ICT01-bound BTN2A1/BTN3A1/BTN3A2 complex reveals significant steric hindrance between the TCR and BTN molecules.

**Supplementary Table S1.**
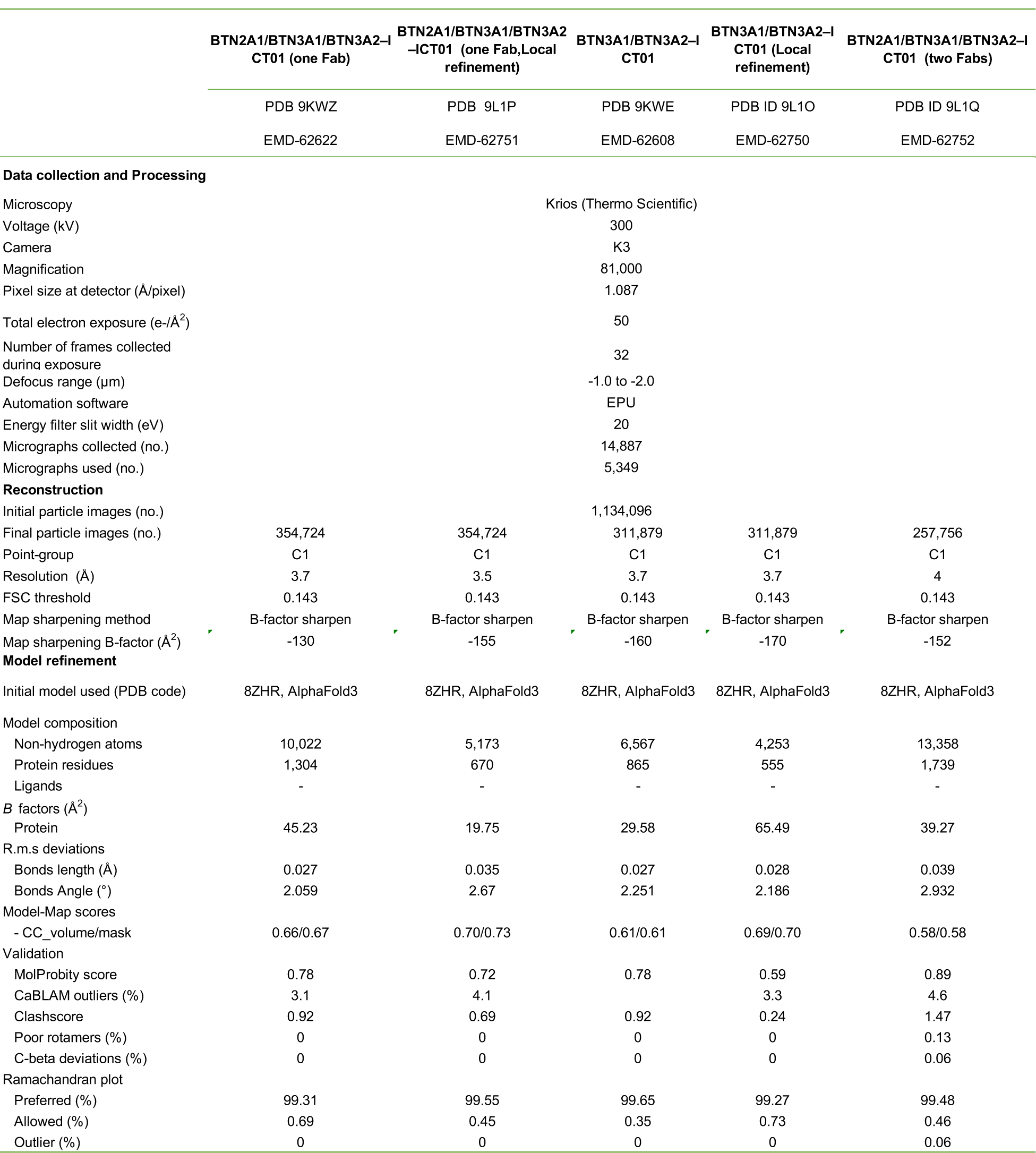
Cryo-EM data collection, refinement and validation statistics.

